# High-throughput and dosage-controlled intracellular delivery of large cargos by an acoustic-electric micro-vortices platform

**DOI:** 10.1101/2021.02.16.431546

**Authors:** Mohammad Aghaamoo, Yu-Hsi Chen, Xuan Li, Neha Garg, Ruoyu Jiang, Abraham P. Lee

## Abstract

Intracellular delivery of cargos for cell engineering plays a pivotal role in transforming medicine and biomedical discoveries. Recent advances in microfluidics and nanotechnology have opened up new avenues for efficient, safe, and controllable intracellular delivery, as they improve precision down to the single-cell level. Based on this capability, several promising micro- and nanotechnology approaches outperform viral and conventional non-viral techniques in offering dosage-controlled delivery and/or intracellular delivery of large cargos. However, to achieve this level of precision and effectiveness, they are either low in throughput, limited to specific cell types (e.g., adherent vs. suspension cells), or complicated to operate with. To address these challenges, here we introduce a versatile and simple-to-use intracellular delivery microfluidic platform, termed Acoustic-Electric Shear Orbiting Poration (AESOP). Hundreds of acoustic microstreaming vortices form the production line of the AESOP platform, wherein hundreds of thousands of cells are trapped, permeabilized, and mixed with exogenous cargos. Using AESOP, we show intracellular delivery of a wide range of molecules (from <1 kDa to 2 MDa) with high efficiency, cell viability, and dosage-controlled capability into both suspension and adherent cells and demonstrate throughput at 1 million cells/min per single chip. In addition, we demonstrate AESOP for two gene editing applications that require delivery of large plasmids: i) eGFP plasmid (6.1 kbp) transfection, and ii) CRISPR-Cas9-mediated gene knockout using a 9.3 kbp plasmid DNA encoding Cas9 protein and sgRNA. Compared to alternative platforms, AESOP not only offers dosage-controlled intracellular delivery of large plasmids (>6kbp) with viabilities over 80% and comparable delivery efficiencies, but also is an order of magnitude higher in throughput, compatible with both adherent and suspension cell lines, and simple to operate.

## INTRODUCTION

Intracellular delivery is an important yet challenging step in gene and cell-based therapies (*1–3*), biomanufacturing (*4, 5*), and basic research (e.g. cell biology, drug discovery, and genetics). Viral vectors are the most widely intracellular delivery method adopted in clinical applications due to their high efficiency and specificity. However, key challenges remain in terms of cytotoxicity, immunogenicity, risk of insertional toxicity, manufacturing, and limited packaging capacity (*6*). Cationic lipids and polymers are among attractive non-viral candidates to replace viral methods, as they cause lower adverse immune responses and have the potential for low-cost and large-scale production. Nevertheless, low delivery efficiency for suspension cells and concerns over cytotoxicity are two major obstacles for these synthetic vectors (*7, 8*). Bulk electroporation is another popular non-viral method for intracellular delivery. Despite its success in delivery of wide range of cargos into most types of cell, including hard-to-transfect cells, high cell mortality is still a major challenge (*9*). In addition, due to their bulk nature, cationic lipids/polymers and electroporation do not offer uniform and dosage-controlled delivery across cell population (*10*).

To address the challenges facing viral and conventional non-viral techniques, microfluidics and nanotechnology have appeared as powerful tools that have shown tremendous potential for adoption in clinical settings and research labs (*11*). Notable examples include methods based on cell deformation (*12, 13*), nanostructures for localized electroporation (*10, 14–16*), mechanoporation (*17, 18*), acoustofluidics sonoporation (*19*), flow-through electroporation (*20*), droplet microfluidics (*21*), and inertial microfluidics (*18, 22*). For safe, efficient, and controllable intracellular delivery, these methods focus on precise control of cellular permeabilization and uptake, down to the single-cell level. To achieve this, cells are usually treated in a 1D or 2D manner. 1D methods flow cells one-by-one and/or usually have channel dimensions at the scale of single cells (*12, 18–21*), while 2D methods are based on monolayer cell culturing or cell interaction with a substrate (*10, 14–17*). Particularly, several existing micro- and nanotechnology methods have adopted such strategies to outperform viral and conventional non-viral techniques in (i) dosage-controlled delivery (*10, 22–26*), which enables the cell population to receive the right concentration of cargo and, thus, minimizes overdose and underdose intracellular delivery, and (ii) intracellular delivery of large cargos (*13, 15, 27–29*), which plays a key role in many genome-editing approaches such as those using CRISPR-Cas9 technology. However, these methods are either low in throughput, limited to specific cell types (e.g., adherent vs. suspension cells), or complicated to operate with.

To address these limitations, here, we present an Acoustic-Electrical Shear Orbiting Poration (AESOP) platform for intracellular delivery of a wide range of cargos with high efficiency, uniformity, cell viability, and throughput of 1 million cells/min per single chip. Compared to existing methods that offer intracellular delivery of large cargos and/or dosage-controlled capability, AESOP is an order of magnitude higher in throughput, compatible with both adherent and suspension cell types, and simple to operate. AESOP incorporates our lateral cavity acoustic transducer (LCAT) technology assisted by interdigitated array (IDA) electrodes for intracellular delivery (Fig. 1 A to C and Movie S1). Once the cells are introduced into the platform, they are consecutively trapped in the array of acoustic microstreaming vortices generated by the LCATs. Since delivery cargos are also pumped along with the cells, they uniformly mix with cells trapped in the microvortices. We hypothesize 3 underlying principles of our AESOP platform: (1) cells trapped in acoustic microstreaming vortices experience modest and uniform mechanical shear that opens small pores on their cell membranes; shear-induced cell membrane poration facilitates intracellular delivery of small molecules (<10KDa) into cells, is cell-dependent, and can be controlled by tuning acoustic transducers, (2) rapid tumbling of cells in the streaming orbits expose them to uniform strength electric fields that uniformly enlarges the pre-existing pores; for formation of large pores, our two-step membrane permeabilization strategy only requires gentle and low-strength electric fields that, alone, are not effective in the absence of microvortices, and (3) vortices induce chaotic mixing, enabling uniform, dosage-controlled and rapid delivery of exogenous materials into the cells.

**Fig. 1.**
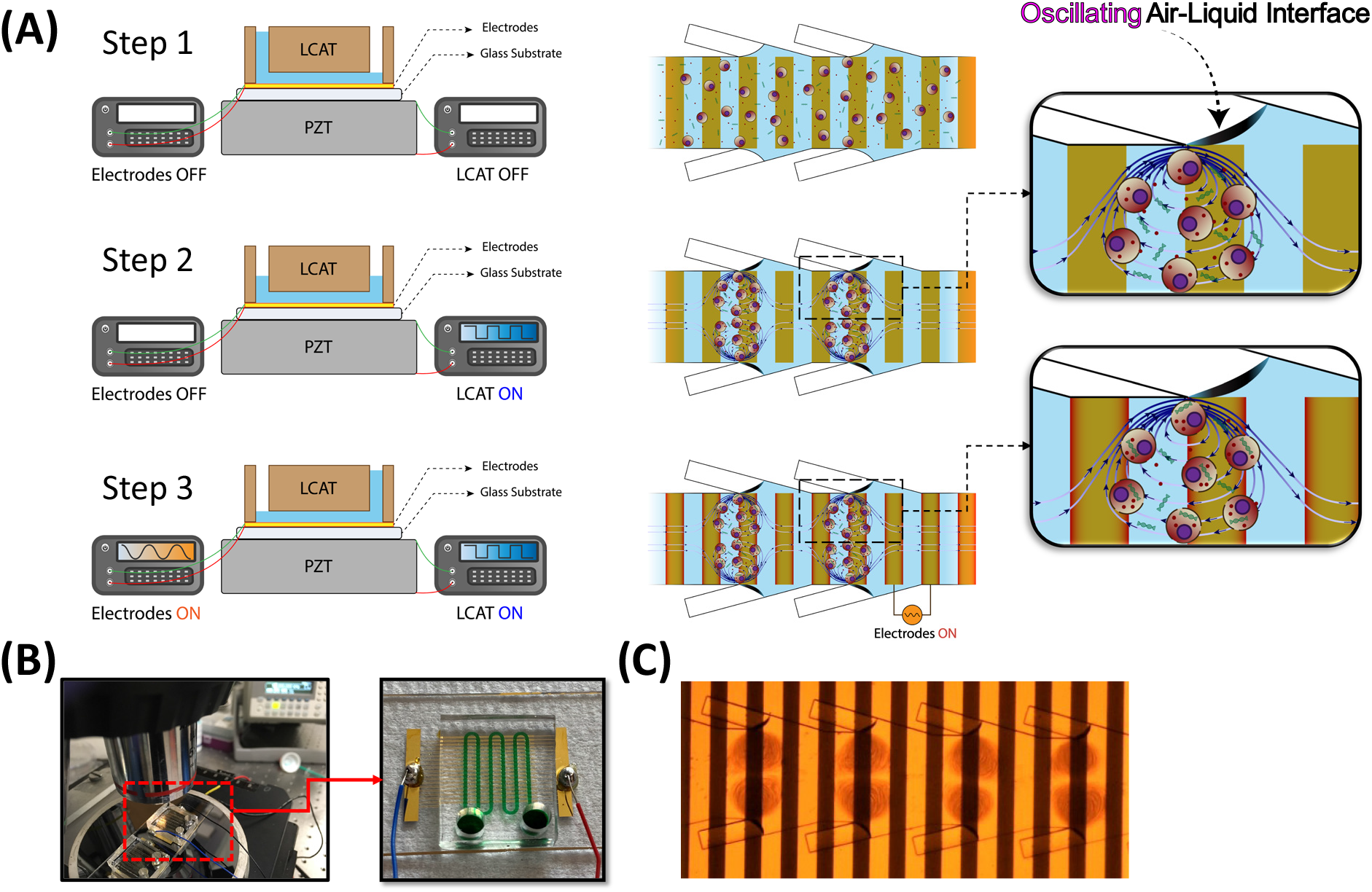
Design and operation of AESOP. (**A**) AESOP’s operational principle is based on three steps: (1) loading the cells and cargos into the chip. Once the solution primes the main channel, air-liquid interfaces will be formed between the main and side channels. (2) Turning on LCAT by applying a resonating square-wave signal to a piezoelectric transducer underneath the chip. The acoustic wave, transmitted from the piezoelectric transducer to the chip, oscillates the air-liquid interfaces, resulting in formation of acoustic microstreaming vortices. The cells trapped in these vortices experience modest and uniform mechanical shear that creates small pores on their membrane. (3) Uniform enlargement of pores by uniform exposure of the cells, rotating in vortices, to the electric field. The cargos are uniformly delivered into cells by chaotic mixing generated by acoustic microstreaming vortices, (**B**) AESOP device setup, and (**C**) Microscope image of cells rotating in acoustic microstreaming vortices, on top of electrodes.

We tested the performance of AESOP with different sizes of molecules, ranging from <1 kDa to 2 MDa, and obtained >90% delivery efficiency with >80% cell viability for both adherent and suspension cell lines. To evaluate AESOP performance for intracellular delivery of large cargos, we first picked a 6.1 kbp enhanced green fluorescent protein (eGFP)-expressing plasmid, and could achieve >80%, >50%, and >40% transfection efficiency for HeLa, K562, and Jurkat cells, respectively, while maintaining high cell viability of >80% for all these cell lines. Using AESOP platform, CRISPR-Cas9-mediated gene editing was demonstrated by a 9.3 kbp plasmid DNA encoding Cas9 and sgRNA to knockout PTEN gene in K562 cells. We showed >80% intracellular delivery of CRISPR plasmid and up to 20% gene knockout across cell population. The large size of the plasmid DNA for eGFP transfection (i.e. 6.1 kbp) and CRISPR-Cas9 gene knockout (i.e. 9.3 kbp) were chosen to challenge the packaging limit of some of the common viral vectors including adeno-associated viruses (AAVs) (*30*). In AESOP, dosage-controlled delivery capability is achieved by the acoustic microstreaming vortices in the key steps of intracellular delivery: (i) membrane disruption: by uniform exposure to mechanical shear and electric field, and (ii) cellular uptake: by uniform mixing of cells with exogenous materials. Delivery analysis of YOYO 1-labelled plasmid DNA confirmed uniform and controllable intracellular delivery across cell population.

Compared to existing methods, our system not only can deliver a wide range of molecular sizes at high efficiency, viability, and uniformity, but it also offers unique sample processing advantages. For example, the unique design of LCATs generates a bulk flow that eliminates the need and complexity of external pumping. In addition, since cells are trapped and suspended in microstreaming vortices, the microfluidic channels are wider, making them higher throughput and less prone to clogging. Furthermore, we have demonstrated a single-chip AESOP platform at a relatively high throughput of up to 1 million cells/min per single chip. This scalability in throughput is relatively straightforward without reduction in system performance.

## RESULTS

### Shear-induced cell membrane poration by acoustic microstreaming vortices

To eliminate the need for applying high electric fields for intracellular delivery, AESOP initiates nanopores on the cell membrane by mechanical shear and enlarges the pre-existing nanopores at lower electric field strengths. To achieve this, AESOP incorporates LCAT technology to trap cells inside acoustic microstreaming vortices and uniformly expose them to modest mechanical shear. The basic structure design of LCAT is illustrated in Figure 2A consisting of a main fluid channel with slanted dead-end side channels. Once the main channel is primed with the sample, air-liquid interfaces are formed along the channel length. When placed on a piezoelectric transducer (PZT), the acoustic energy is transmitted to the air-liquid interfaces of LCATs, causing them to oscillate and generate microstreaming vortices in the microfluidic channel. The orientation and positioning of the air-liquid cavities result in both bulk flow liquid pumping and size-selective trapping of cells (*31, 32*). The trapped cells orbiting in these micro-vortices are subjected to oscillatory mechanical shear, which can be controlled by varying the interface oscillation amplitude excited by the PZT.

**Fig. 2.**
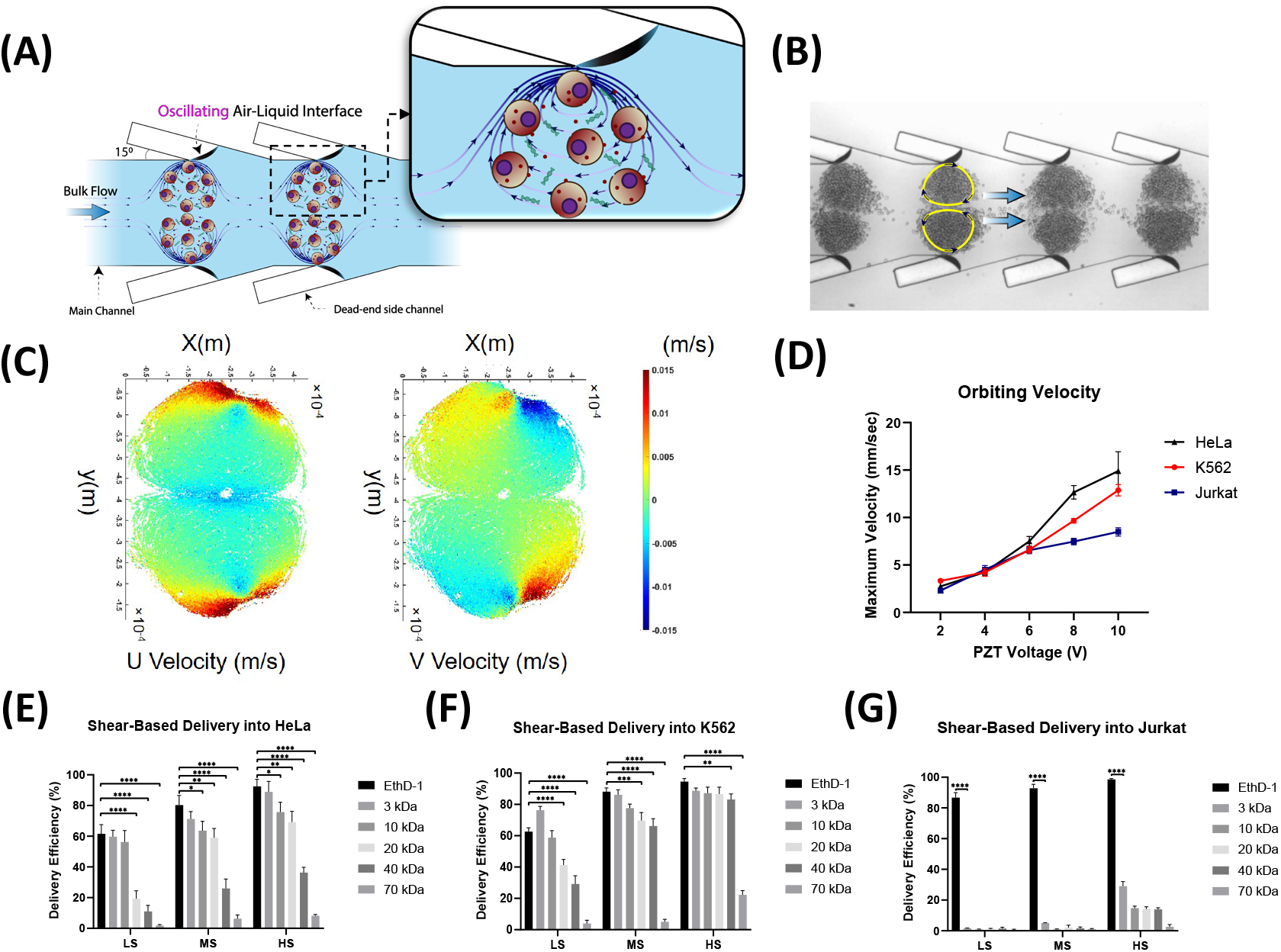
Shear-induced initiation of small pores on cell membrane by acoustic microstreaming vortices. (**A**) Basic design structure of LCAT technology incorporated by AESOP for shear-induced small pore formation; LCATs are arrays of acoustically actuated air-liquid interfaces generated using dead-end side channels, (**B**) Microscope image of K562 cells trapped in acoustic microstreaming vortices generated by LCAT, (**C**) PTV analysis results of K562 cells orbiting in acoustic microstreaming vortices (PZT voltage= 6V), (**D**) Cells’ maximum velocity orbiting in micro-vortices; the maximum velocity is reached near the air-liquid interface, and is proportional to the PZT applied voltage, (**E**-**G**) Shear-induced delivery of small molecules into (**E**) HeLa, (**F**) K562, and (**G**) Jurkat cells at three different operational modes: “low shear (LS)” (PZT voltage = 2V), “moderate shear (MS)” (PZT voltage = 6V), and “high shear (HS)” (PZT voltage = 10V). *P<0.05, **P<0.01, ***P<0.001, and ****P<0.0001 were determined by Tukey’s honest significant difference criterion.

In the first step, particle tracking velocimetry (PTV) was employed to characterize micro-vortices and measure the velocity and trajectory of cells at different PZT applied voltages (Fig. 2, C and D, and *SI Appendix,* Fig. S1-S3). The PTV results indicate that the cells’ maximum velocity is reached near the air-liquid interface (Fig. 2C), and its magnitude is directly proportional to the PZT applied voltage (Fig. 2D). In addition, by increasing the PZT applied voltage, the device pumping rate increases linearly (*SI Appendix,* Fig. S4).

In the next step, we tested the hypothesis of using acoustic microstreaming vortices for shear-induced initiation of nanopores on cell membrane. For this purpose, we introduced cells (HeLa, K562, or Jurkat) with different molecules (EthD-1 dye (~857 Da) or fluorescein isothiocyanate–dextran with different sizes, ranging from 3 kDa to 70 kDa) into the chip and activated the LCAT. To evaluate the effect of mechanical shear force on intracellular delivery, we picked three different PZT applied voltages corresponding to “low shear (LS)” (PZT voltage = 2V), “moderate shear (MS)” (PZT voltage = 6V), and “high shear (HS)” (PZT voltage = 10V) (Movie S2). Based on the results (Fig. 2, E to G), there are four key findings: 1) Mechanical shear facilitates delivery of small molecules into the cells, indicating formation of nano-sized pores on the cells’ membrane, 2) At a given shear rate, the delivery efficiency of larger cargos is lower than smaller cargos, 3) Increasing the shear increases the delivery efficiency of molecules into the cells by creating larger pores, 4) there exists a pore size threshold for shear-induced cell membrane poration; for the three shear modes (low, moderate, and high) tested, shear alone could not deliver >1kDa molecules into Jurkat cells and ≥70kDa molecules into HeLa and K562. Based on these results, the size of generated pores is mainly dependent on PZT applied voltage and cell type. Even though high shear mode provides higher delivery efficiency, for the rest of studies, we chose moderate shear (PZT voltage = 6V) as our optimum operational mode for LCAT. This mode offers effective small pore formation (>80% delivery efficiency for molecules up to 3 kDa in size for K562 and HeLa, and >90% delivery efficiency of ~857 Da EthD-1 dye for Jurkat). Importantly, compared to high shear mode, moderate shear mode is gentler on the cells, especially when coupled with electric field pore enlargement modality.

### Uniform electric field enlargement of shear-induced pores for cargo delivery

Once small pores on cells’ membrane are initiated by acoustic microstreaming vortices, AESOP enlarges the pores by applying a sinusoidal AC electric field via interdigitated array (IDA) electrodes. For each different cell type, electric field voltage, frequency, and applied time were optimized (*SI Appendix*, Supplementary Note 1, and Fig. S5-S7). Specifically, we found 12.5 V_max_, 10 kHz, and 10 ms for HeLa cells, 35 V_max_, 30 kHz, and 10 ms for K562 cells, and 25 V_max_, 20 kHz, and 10 ms for Jurkat cells, as the optimum electric field parameters. We, then, tested the performance of AESOP in delivery of dextran with molecular sizes ranging from 3 kDa to 2 MDa (Fig. 3A). According to the results, for all three different cell lines tested, >90% delivery efficiency was achieved for any given molecular size of dextran.

**Fig. 3.**
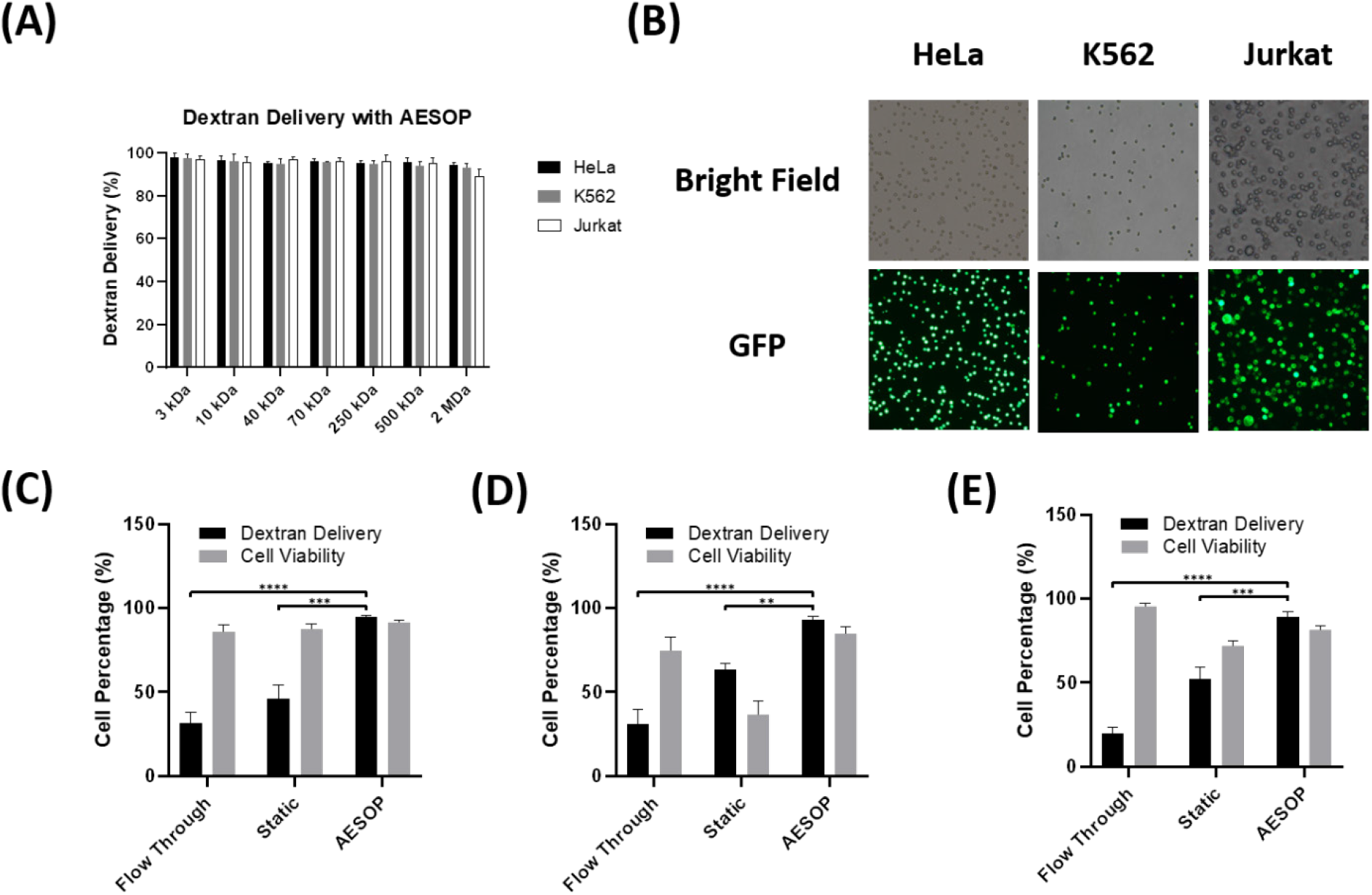
AESOP for intracellular delivery of different sizes of cargo. (**A**) Performance of AESOP in delivery of dextran, with wide range of molecular size, into HeLa, K562, and Jurkat cell lines, (**B**) Brightfield and GFP image of HeLa, K562, and Jurkat cells tested for delivery of 2 MDa dextran with AESOP platform, (**C**-**E**) Comparison of the performance of the AESOP with “Flow Through” and “Static” in delivery of 2 MDa dextran and corresponding cell viability for (**C**) HeLa, (**D**) K562, and (**E**) Jurkat; the results indicate that AESOP offers significantly higher delivery efficiency compared to the “Flow through” and “Static” groups. *P<0.05, **P<0.01, ***P<0.001, and ****P<0.0001 were determined by Tukey’s honest significant difference criterion.

In the next step, to evaluate the role of shear in AESOP performance, we fixed the optimum applied electric field parameters for each cell type and compared AESOP with: 1) Flow-through: with LCAT off, the cells were flown through the chip and on top of the electrodes using a syringe pump, and 2) Static: with LCAT off, the cells were loaded into the chip and settled down on top of the electrodes. The results for delivery of 2 MDa dextran (Fig. 3, C to E) indicate that AESOP achieves higher delivery efficiency (>90%) compared to flow-through (low, <30%) and static (moderate, <60%). The moderate delivery efficiency of static approach is because the cells are close to or in contact with the electrodes and, thus, experience high electric field strengths. Like bulk electroporation, the exposure to high electric field in static approach is particularly unfavorable for applications that long-term cell viability is critical after the delivery process (e.g. plasmid and mRNA delivery).

### Dosage-controlled capability and mechanism of intracellular delivery

In AESOP platform, acoustic microstreaming vortices play a key role in efficient and precise intracellular delivery of cargos. The cells in these vortices are not only exposed to uniform mechanical shear and electric field, but also uniformly mixed with exogenous cargos by chaotic mixing. Thus, we hypothesized that the imposed uniformity in membrane disruption and cellular uptake would result in dosage-controlled intracellular delivery across cell population. To test this hypothesis, we delivered YOYO-1 labeled plasmid DNA (6.1 kbp) into K562 cells using AESOP and two other control groups (static and flow through). Since the amount of plasmid DNA delivered to a cell is directly proportional to the measured fluorescent intensity of YOYO-1 dye within the cell, fluorescent intensity distribution among cell population was analyzed by flow cytometry. According to the histogram of fluorescent intensity (Fig. 4A), AESOP offers a narrow peak distribution of YOYO-1 labeled DNA, indicating delivery of uniform doses across the cell population. In contrast, for the two control groups, where the effect of vortices was eliminated by turning LCAT off, the intensity peak distribution of delivered DNA is wide and not uniform among population of cells. To better quantify controllable intracellular delivery, we calculated the percentage coefficient of variation (%CV, defined as the percentage ratio of standard deviation to the mean) of fluorescent intensity across cell populations processed by the control groups and AESOP with DNA concentrations of 50 ng, 500 ng, 1 μg, and 2.5 μg per million cells (Fig. 4B). Unlike the two control groups with %CV>120, all AESOP groups offer %CV around 50%. The low %CV achieved by AESOP groups not only confirms delivery of uniform doses across the cell population, but also is an indicator of performance consistency when working with different cargo concentrations. In addition, for each different DNA concentrations, we calculated the mean fluorescent intensity of YOYO-1 dye delivered to the cells (Fig. 4C). Based on the results, the average dose delivered to the cells is linearly proportional to DNA concentration, indicating that AESOP offers controllable intracellular delivery. We also investigated whether high-throughput cell processing affects the performance of AESOP for gene delivery. For this, two different AESOP designs were tested: 1) moderate throughput: capable of processing up to 200K cells/min (Fig. 1B and C), and 2) high throughput: capable of processing up to 1M cells/min (*SI Appendix,* Fig. S8, and Movie S3). Based on the results (Fig. 4D and E), except overdose delivery into a small percentage (~5%) of cells in the high-throughput version, there is no significant difference in delivery efficiency and uniformity between the two versions. This indicates that scalability of the platform is straightforward and does not significantly affect system performance. Figure 4F shows the corresponding efficiency and cell viability for delivery of labeled plasmid into cells. Similar to the trend observed in intracellular delivery of 2 MDa dextran, the results indicate that both moderate and high-throughput AESOP versions achieve high delivery efficiency of plasmid (>80%) while static and flow through result in <60% and <30% efficiency, respectively.

**Fig. 4.**
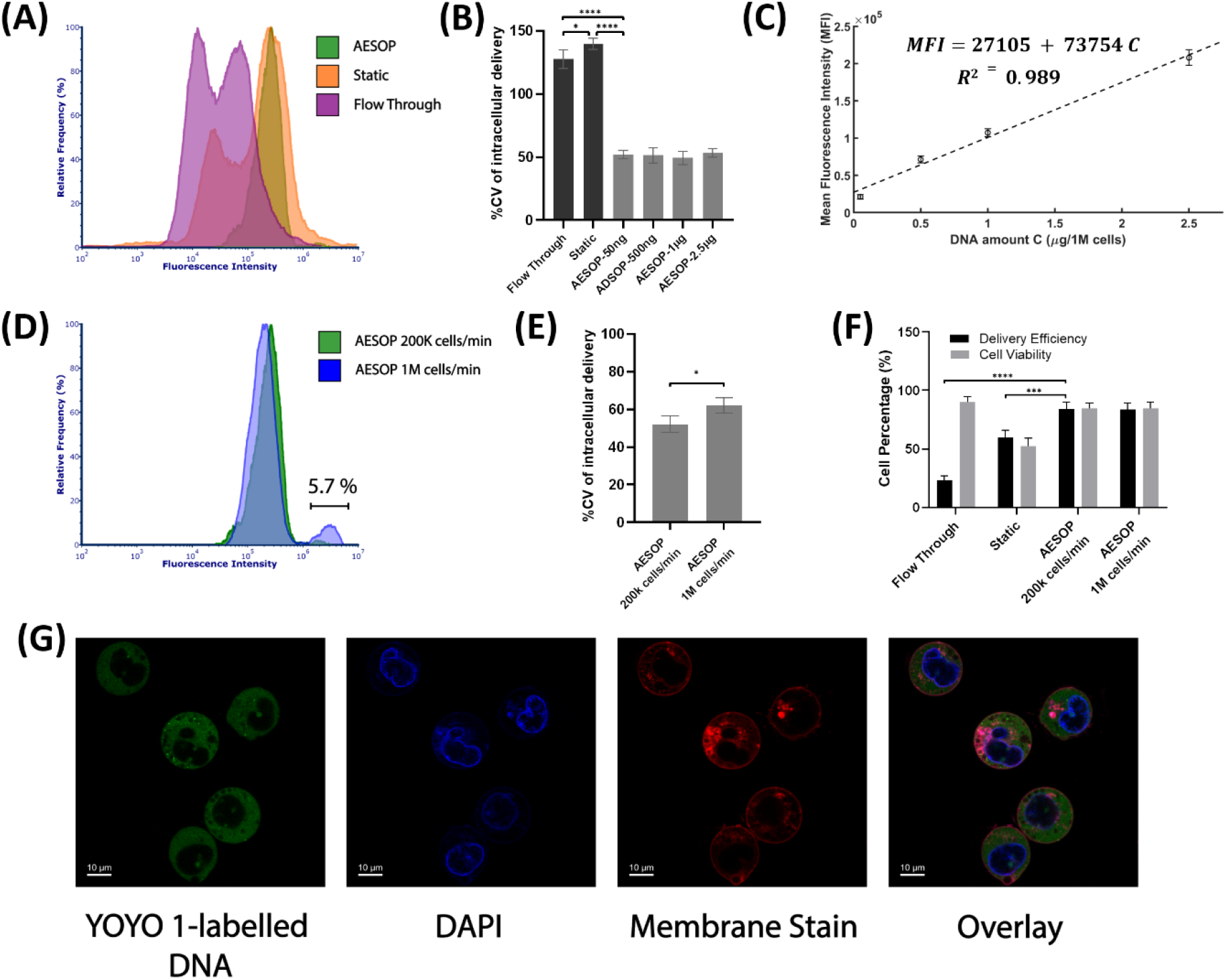
Intracellular delivery of fluorescent labeled plasmid DNA. (**A**) Histogram of fluorescent intensity of YOYO-1 labeled plasmid DNA delivered into K562 cells. For better comparison, the histograms were normalized by relative frequency (%). Compared to control groups, AESOP offers a sharp and narrow intensity distribution, indicating AESOP’s capability for precise and controlled delivery, (**B**) %CV of intracellular delivery for control groups (flow through and static) and AESOP operated with different DNA concentration, (**C**) Mean fluorescent intensity of YOYO-1 dye delivered to the cells by AESOP operated with different DNA concentrations. A linear model is fitted to the obtained data, indicating controlled delivery by AESOP, (**D**) The histogram of fluorescent intensity of YOYO-1 labeled plasmid DNA delivered into K562 cells using moderate and high throughput AESOP platforms, (**E**) %CV of intracellular delivery for moderate and high-throughput AESOP versions; increasing the throughput resulted in slight increase in delivery distribution across cell population, (**F**) The Delivery efficiency and cell viability for intracellular delivery of labeled DNA into cells; the results show that >80% plasmid DNA delivery efficiency can be achieved using AESOP platform, and (**G**) Confocal microscopy image of cells after intracellular delivery experiment with AESOP; for this experiment, the cells’ nucleus and membrane were stained with DAPI and deep red CellMask™ plasma membrane stains. Based on the results, acoustic microstreaming vortices directly deliver the plasmid DNA to the cell cytoplasm. *P<0.05, **P<0.01, ***P<0.001, and ****P<0.0001 were determined by Tukey’s honest significant difference criterion.

In the next step, we evaluated how AESOP facilitated intracellular delivery of plasmid DNA into cells. The main question was whether LCAT entangles the plasmid DNA to the cell membrane (*33*), or it delivers directly to either the cytoplasm or nucleus (*14, 34*). For this purpose, cell nuclei and membranes were labeled with DAPI and CellMask™ plasma membrane stains, respectively. After the experiment, confocal microscopy was performed to observe the distribution of labeled plasmid DNA in K562 cells. Based on the results (Fig. 4G), the plasmid DNA are mostly delivered into the cells’ cytoplasm. Since AESOP utilizes AC electric field with frequencies ≥10kHz, the effect DNA electrophoresis can be neglected. This indicates that the chaotic mixing induced by microstreaming vortices acts as the major active force to guide the plasmid DNA through the cell membrane, and into the cytoplasm. As a result, LCAT eliminates the need for any other active force, such as electrophoresis, to guide DNA into the cells.

### Gene delivery analysis: eGFP plasmid DNA transfection & CRISPR-Cas9 gene editing

We also explored the performance of AESOP for intracellular gene delivery applications in protein expression and targeted gene knockout. First, a relatively large sized eGFP-expressing plasmid DNA (6.1 kbp) was chosen and delivered into HeLa, Jurkat, and K562 cells using the AESOP platform. Flow cytometry results 48 hrs after the gene delivery experiment were analyzed and GFP protein expression efficiencies of >80%, >40%, and >50% were obtained for HeLa, Jurkat, and K562 cells (Fig. 5, A and B). In addition, high cell viability of >80% was achieved for all the three cell types tested. We then evaluated AESOP for CRISPR-Cas9 gene editing applications. For this purpose, a 9.3 kbp plasmid DNA encoding Cas9 protein and sgRNA targeting PTEN gene knockout were chosen and delivered into K562 cells. Based on the flow cytometry analysis of cells treated with AESOP, CRISPR-plasmid intracellular delivery efficiencies of >80% were achieved for K562 cells (Fig. 5C). After gene delivery, the cells were cultured for 48hrs, selected with eGFP marker, cultured for an additional 7 days, and analyzed by immunofluorescence (IF) staining. Compared to the control group, where PTEN proteins were detected in the cytoplasm, the experimental group showed clear knockout of the gene (Fig. 5D). Overall, we could achieve up to 20% gene knockout via AESOP platform.

**Fig. 5.**
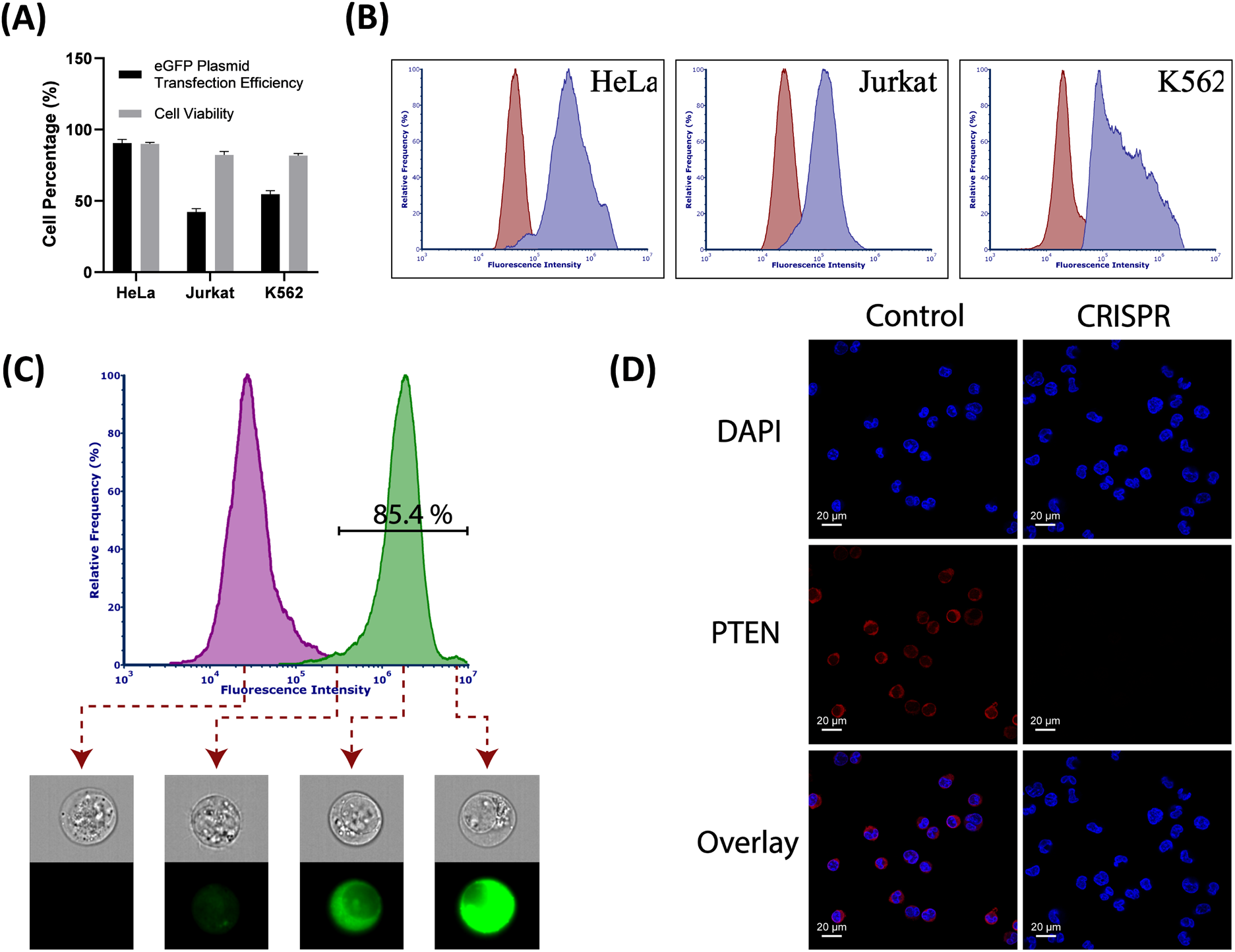
GFP protein expression analysis. (**A**) Transfection efficiencies and cell viability 48 hrs after delivery of 6.1 kbp eGFP-expressing plasmid DNA; AESOP achieved >80%, >40%, and >50% transfection efficiencies with high cell viabilities, (**B**) Flow cytometry quantification of eGFP expression for experimental (blue) and control (red) groups, (**C**) Flow cytometry quantification of delivery of YOYO-1 labelled plasmid DNA encoding Cas protein and PTEN sgRNA into K562 cells for experimental (green) and control (purple) groups. For better comparison, the histograms were normalized by relative frequency (%), (**D**) immunofluorescence (IF) staining of K562 cells with PTEN monoclonal antibody recognized by Alexa Fluor 647 secondary antibody. DAPI was used to stain the nucleus.

## DISCUSSION

AESOP is a multimodal non-viral intracellular delivery platform that meets the key criteria needed for adoption in gene/cell-based therapies, biomanufacturing, and basic research. These criteria include: i) High delivery efficiency while maintaining cell viability, ii) Dosage-controlled delivery of cargos, iii) High throughput, iv) Compatibility with both adherent and suspension cell types, and v) Simplicity in fabrication and operational protocol. For this, AESOP controls cell membrane permeabilization and cellular uptake in an efficient, precise, and high-throughput manner.

To permeabilize cell membranes effectively and gently, AESOP adopts a two-step membrane disruption strategy. First, it forms nanopores on the cell membrane using mechanical shear. Second, it enlarges these nanopores upon the cells’ uniform exposure to gentle electric fields. Stable bubble oscillations has been known to apply local shear force and permeabilize nearby cells via sonoporation (*35, 36*). Similar to this principle, AESOP employs LCAT’s acoustic microstreaming vortices to apply tuned and moderate mechanical shear on cells and, consequently, create small pores on their membrane. To open larger pores, AESOP needs only gentle and low-strength, rather than undesirable high-strength, electric fields. This strategy is indeed similar to dual-pulse electroporation strategy where the cells experience a short, high-strength pulse followed by a long, low-strength pulse. The former creates several small pores on cell membrane, and the latter expands the pores and electrophoretically guides the charged cargos into the cell (*37, 38*). This strategy has shown to improve delivery efficiency and cell viability (*11*). AESOP outperforms dual-pulse electroporation technique because it does not rely on high-strength pulse in any steps. As a result, it overcomes fundamental challenges of using high-strength electric fields in electroporation such as joule heating, metal contamination, electrolysis, and pH change in buffer. In addition, since cells are tumbling in acoustic microstreaming vortices, they are uniformly exposed to both mechanical shear and electric field, resulting in uniform membrane permeabilization across cell populations.

In terms of cellular uptake, AESOP uses chaotic mixing, induced by acoustic microstreaming vortices, to deliver cargos efficiently and uniformly into cells. The majority of intracellular delivery approaches rely on either passive diffusion or electrophoresis to guide the cargos into cells. Compared to passive diffusion, electrophoresis significantly improves intracellular delivery of cargos into permeabilized cells. Recently, micro-vortices have also shown to enhance cellular uptake by mixing cells with the exogenous cargos (*20–22*). In our previous work, we developed a droplet microfluidic platform for lipid-mediated single-cell transfection. We showed that chaotic advection, formed inside droplets moving in a winding channel, can significantly enhance cell transfection efficiency and uniformity (*21*). Here, in AESOP, we took a new approach for high-throughput and efficient mixing, and designed hundreds of whirlpool-like microstreaming vortices to simultaneously mix hundreds of thousands of permeabilized cells with exogenous cargos. Our presented results indicate the important role of these vortices to increase cellular uptake efficiency.

As a result of uniform membrane permeabilization and cellular uptake, AESOP offers dosage-controlled delivery capability. This is an important requirement for many cell engineering applications. For example, Mali et al. showed that precise control over Cas9-sgRNA dose is critical for achieving desired targeting specificity in Cas9 gene editing (*39*). In this paper, we evaluated dosage-controlled intracellular delivery by flow cytometry analysis of the cells processed by AESOP. We used %CV as an indicator of relative dispersion of the amount of DNA delivered to the cell population and showed that %CV<60 can be achieved by AESOP. In the next step, AESOP performance was evaluated at different DNA concentrations. We found out that: (i) %CV is independent of the cargo concentration, showing the performance consistency, and (ii) the average dose delivered to individual cells is linearly proportional to the cargo concentration. Utilizing this precise intracellular delivery approach, we could lower the cargo concentration, down to 1 μg of plasmid per million cells. This is particularly important for reducing the cost and minimizing toxicity associated with plasmids (*40*). Overall, several promising micro- and nanotechnology approaches have been developed for dosage-controlled intracellular delivery such as nanostraw-electroporation (*10*), nanofountain probe electroporation (*23*), nanochannel electroporation (*24*), micro/nano-injection (*25, 26*), and microscale symmetrical electroporator arrays (*22*). Compared to these methods, AESOP is an order of magnitude higher in throughput and is not cell type-dependent, as cells are suspended in acoustic microstreaming vortices.

In recent years, there has also been growing need for intracellular delivery of large cargos for gene editing. For example, most plasmid-based CRISPR-Cas9 gene editing cargos are >9 kbp. Another powerful recent development is base editing, which uses a cytosine base editor (CBE) or an adenine base editor (ABE) with a guide RNA with an approximate size of 6 kbp (*41*). For these applications, the use of viral vectors is challenging due to their limited packaging capacity. For example, AAVs, as popular vectors for gene/cell-based therapies, have a packaging capacity of 4.7 kbp and, thus, dual or triple AAV delivery approaches are required for cargos that exceed such a limit (*41, 42*). Recently, methods based on membrane deformation (*13*), bubble cavitation (*27, 28*), nanochannel electroporation (*15*), and high-frequency ultrasound (*29*) have demonstrated successful delivery of large cargos. Despite encouraging results, compared to AESOP, these methods are lower in throughput and/or require cell interaction with a substrate that limits their application mostly to adherent cells. Our presented results with 6.1 kbp eGFP plasmid and 9.3 kbp CRISPR-Cas9 plasmid show that AESOP also addresses two other key challenges associated with delivery of large cargos: low delivery efficiency and cell viability. Overall, it is more challenging to deliver larger cargos, as diffusion-limited intracellular delivery becomes extremely inefficient. One solution is to increase the cargo concentration to achieve acceptable transfection efficiency. Thus, for large cargos, not only are larger pores needed, high concentration of cargos greatly reduces the cells’ viability, in particular for plasmid-based gene editing applications as there exists specific toxicity associated with large plasmids (*40*). The chaotic advection provided by AESOP not only increases transport of larger cargo molecules to overcome their lower diffusion rates, but it also reduces the required concentration of cargos to minimize overdose delivery across cell populations.

For adoption in clinical settings, intracellular delivery platforms should also satisfy the requirement for high-throughput cell processing. As an example, Tisagenlecleucel (Kymriah), the anti-CD19 chimeric antigen receptor (CAR) T-cell therapy for pediatric patients with B-cell precursor acute lymphoblastic leukemia (ALL), requires an average dose of 1×10^8^ transduced viable T cells (*43*). Our current 2 cm × 5 cm AESOP chip can already process up to 1 million cells/min. In this work, by comparing moderate (200k cells/min/chip) and high-throughput (1M cells/min/chip) AESOP versions, we demonstrated the scalability of our platform without sacrificing the delivery efficiency and precision across cell populations. This is mainly because AESOP consists of hundreds of microvortices, each holding thousands of cells, that act as independent reactors. With flow control and optimization, we are confident that we can achieve throughput of 100 million cells/min. This would require optimization of the microfluidic channels and LCATs to maximize cell processing density and speed. Increasing the channel dimensions (length, width, height) is the most direct way to increase throughput. If necessary, parallelization of multiple chips (stacking) would be also adopted. One of the intrinsic advantages of the AESOP technology is that the whole system is compact with pumping, trapping, shearing, and interdigitated electrodes, all on one common microfluidic chip platform. Consequently, the whole system is simple, easy to operate, test, and characterize.

## MATERIALS AND METHODS

### Materials and reagents

Dulbecco’s modified Eagle’s medium (DMEM), fetal bovine serum (FBS), Iscove’s modified Dulbecco’s medium (IMDM), Roswell Park Memorial Institute (RPMI) 1640 were purchased from Thermo Fisher Scientific. HeLa, K562, and Jurkat (human acute T cell leukemia cell line) cells were purchased from American Tissue Culture Collection (ATCC; Manassas, VA). Fluorescein isothiocyanate–dextran molecules were purchased from MilliporeSigma. pcDNA3.1+C-eGFP plasmid and plasmid encoding Cas9 and sgRNA were purchased from GenScript^®^. For PTEN targeted gene knockout experiment, the 20bp sgRNA sequence of TTATCCAAACATTATTGCTA was used. YOYO-1 dye (1 mM solution in DMSO; Invitrogen, cat. no. Y3601), DAPI (4’,6-diamidino-2-phenylindole) stain, PTEN Monoclonal Antibody, and Goat anti-Mouse IgG (H+L) Highly Cross-Adsorbed Secondary Antibody, Alexa Fluor Plus 647 were purchased from Thermo Fisher Scientific.

### Device Fabrication

AESOP integrates interdigitated array (IDA) electrodes with LCAT chip (*SI Appendix,* Fig. S9). Lift-off technique was adopted for batch electrode fabrication. For this, the glass slides were, first, cleaned with acetone, isopropyl alcohol, and methanol, and dried overnight at 120°C. Standard photolithography, using MICROPOSIT™ S1813 positive photoresist, was performed to fabricate patterns on the glass slides. Using e-beam evaporation, 300°A chromium (Cr) followed by 1000°A gold (Au) was deposited on the slides. The Cr layer was chosen to improve the adhesion of Au layer to the substrate. After thin film deposition, the glass slides were sonicated in a bath of acetone to remove the photoresist and, subsequently, stripping away unwanted metal layers. Soft lithography technique was employed for fabrication of the LCAT chip. For this, negative photoresist SU-8 2050 (Kayaku Advanced Materials, Inc.) was used for pattern fabrication on silicon wafer. The silicon wafer was then silanized overnight by (TRIDECAFLUORO-1,1,2,2-TETRAHYDROOCTYL)TRICHLOROSILANE (Gelest, Inc.) to avoid PDMS-mold adherence. Poly (dimethylsiloxane), PDMS (Sylgard 184, Dow Corning) base and curing agent were mixed at the ratio of 10:1, poured on the mold, degassed for 1 hour in a desiccator, and cured at 65°C overnight. The cured PDMS was peeled from the wafer, cut to size, and cleaned. Both the LCAT chip and glass slide, with patterned electrodes, were aligned and bonded by oxygen plasma treatment. Finally, to make the AESOP device hydrophobic, it was baked overnight at 65°C.

### Standard AESOP operation and intracellular delivery protocol

For efficient transport of acoustic wave from PZT to the device, an ultrasound gel (Aquasonic) was first smeared between the AESOP chip and the PZT (STEMiNC, STEINER & MARTINS, Inc.). The PZT and AESOP chip were separately connected to a signal generator (Agilent 33220A) and a power amplifier (JUNTEK^®^). The cell solution was suspended in electroporation buffer (Bio-Rad Laboratories, Inc.), mixed with the desired concentration of exogenous cargos (e.g., dextran, eGFP plasmid, CRISPR plasmid, etc.), and pipetted into the device inlet in 30 μL sample batches. For pumping the sample into the chip and applying tunable mechanical shear to the cells, the PZT was then excited by an square wave at fixed frequency of 50.2 kHz and the desired amplitude. For electrical expansion of shear-induced pores on cells’ membrane, a sinusoidal wave was applied to the interdigitated array (IDA) electrodes with the desired frequency, amplitude, and duration. It should be mentioned that throughout device operation, LCATs were turned OFF and ON a few times for better mixing of cells and cargos. After delivery, the cells were collected from the device and recovered in cell culture medium without fetal bovine serum (FBS) for 20 minutes. After recovery, the cells were dispersed in their respective culture media supplemented with 10% FBS and incubated in a humidified atmosphere of 5% CO2/95% air at 37°C.

### Cell culture

HeLa cells were grown in DMEM supplemented with 10% FBS. K562 cells were grown in IMDM supplemented with 10% FBS. Jurkat cells were grown in RPMI 1640 medium supplemented with 10% FBS. All cells were cultured in a humidified atmosphere of 5% CO2/95% air at 37°C.

### Particle Tracking Velocimetry (PTV) analysis

Cells’ motion in acoustic microstreaming vortices was captured using a high-speed camera (Phantom, vision research) connected to a L150 Nikon Eclipse upright microscope. For improved particle detection, high pass filter was used for edge detection. The video was then analysis by an open-source MATLAB code to track the cells and obtain their velocity components in microstreaming vortices (*44*).

### Labeling plasmid DNA, immunofluorescence (IF) staining, flow cytometry, and confocal laser scanning microscopy

For studying mechanism, efficiency, and uniformity of intracellular delivery, plasmid DNA was labeled with YOYO-1 dye at a ratio of 1 dye molecule per 5 bp of the DNA (*20*). For this, the desired concentration of plasmid DNA was mixed with the YOYO-1 dye and incubated for 1 hour at room temperature in the dark. The fluorescently labeled DNA was then mixed with the cell sample and intracellular delivery was performed. After sample collection, the cells were washed three times in 1X PBS to remove background and any nonspecifically adsorbed plasmid DNA from the cell surface. The cells were then resuspended in 1X PBS for flow cytometry and confocal laser scanning microscopy.

IF analysis was adopted to evaluate the CRISPR-Cas9–mediated targeted gene knockout efficiency. For this, the cells were first fixed using 4% formaldehyde (pH 7.4) (Polysciences, Inc.) for 10 minutes at room temperature. Then, they were permeabilized by 0.1% Triton X-100 (ICN Biomedicals, Inc.) in 1X PBS for 15 minutes at room temperature and blocked with 3% BSA in PBST (PBS+ 0.1% Tween 20) for 30 minutes at room temperature. Cells were then probed with the diluted primary PTEN antibody in 1% BSA in PBST (1:10 dilution ratio) overnight at 4°C. They were then incubated with the secondary antibody (5 μg/ml) in 1% BSA for 1 hour at room temperature in the dark. The cells were then resuspended in 1X PBS and plated on a microscope slide for confocal laser scanning microscopy. In between all the IF steps, the cells were washed in 1X PBS three times.

Flow cytometry was performed by an ImageStream Mark II Imaging Flow Cytometer (Amnis Corporation) at 60× magnification under the laser excitation of 488 nm. Confocal laser scanning microscopy was performed by a ZEISS LSM 700 laser scanning confocal microscope (Carl Zeiss) with a 63x oil objective and three laser lines: 405nm for DAPI, 488nm for YOYO-1 labelled plasmid, and 639nm for detecting Alexa Fluor Plus 647 secondary antibodies.

### Cell viability test

The Calcein Red AM (AAT Bioquest) was used to determine the cell viability. For analysis, the cells were resuspended in 1x PBS buffer (ThermoFisher Scientific) and stock solution of Calcein Red AM was added to the cell’s solution with 1:100 volume ratio. The flow cytometry was used to evaluate the viability.

### Statistical analysis

One-way ANOVA and multiple comparison with Tukey’s honest significant difference criterion were performed by MATLAB. The percentage coefficient of variation (%CV) of fluorescent intensity of YOYO-1 dye delivered to the cell population were calculated on at least 95% of cell population as:

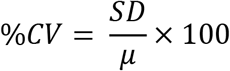

where SD and μ are standard deviation and mean of fluorescent intensity, respectively.

## ACKNOWLEDGEMENTS

The authors would like to acknowledge support from the National Science Foundation and the industrial members of the Center for Advanced Design and Manufacturing of Integrated Microfluidics (NSF I/UCRC award number IIP 1841509).

