## Supplementary Information Appendix for "High-throughput and dosage-controlled intracellular delivery of large cargos by an acoustic-electric micro-vortices platform"

**This PDF file includes:**

Supplementary Note 1

Figures S1 to S9

**Supplementary Note 1: Optimization of electric field parameters for cargo delivery**

AESOP utilizes interdigitated array (IDA) electrodes to enlarge the small pores initiated by acoustic microstreaming vortices. For each different cell type, electric field voltage ($V_{max}$), frequency (f), and applied time (T) were optimized. In terms of starting values, we chose $V_{max}$=7.5V, f=10kHz, and T=10ms for all the cell types. In addition, to reduce the complexity associated with such optimization, we fixed T for the rest of study. The optimization process was based on two steps: (1) Optimizing 2 MDa dextran delivery, and (2) Optimizing eGFP-expressing plasmid DNA (6.1 kbp) transfection efficiency. The role of step 1 was to save time in narrowing down the feasible domains for $V_{max}$ and f. This is mainly because dextran delivery efficiency can be evaluated in a matter of hours while it takes ~48 hours to determine eGFP transfection efficiency. For both steps, cell viability was also considered as an optimization constraint to be >80%.

For HeLa cells, at f=10kHz, we found out that $V_{max}$≥12.5V results in >90% dextran delivery efficiency and acceptable cell viability for $V_{max}$≤17.5V (Fig. S5A). Thus, we narrowed down the applied voltage domain to 7.5V≤$V_{max}$≤17.5V and performed eGFP transfection experiments. Based on the results (Fig. S5B), we obtained $V_{max}$=12.5 as the optimum applied voltage, at which >80% transfection efficiency and cell viability were achieved.

Following the same optimization protocol for Jurkat cells, we could not achieve any desirable transfection efficiency without sacrificing cell viability. As a result, we increased the applied frequency to f=20kHz, where the AC electric field is gentler to the cells. This enabled us to increase the applied voltage while maintaining viability. Based on the dextran results (Fig. S6A), we narrowed down the applied voltage domain to 10V≤$V_{max}$≤30V for eGFP transfection optimization. According to the results (Fig. S6B), we found $V_{max}$=25V as optimum applied voltage that resulted in >40% transfection efficiency and >80% cell viability.

For K562 cells, applied frequencies of 10kHz and 20kHz did not result in high transfection efficiency while maintaining viability >80%. Thus, we increased the applied frequency to 30kHz which, consequently, enabled us to explore higher applied voltages without harming the cells. By dextran experiments, we could narrow down the applied voltage domain to 20V≤$V_{max}$≤40V (Fig. S7A). Exploring this domain (Fig. S7B), we optimized the transfection efficiency with respect to electric field voltage and obtained $V_{max}$=35V as the optimum parameter that gave >50% transfection efficiency and >80% cell viability.

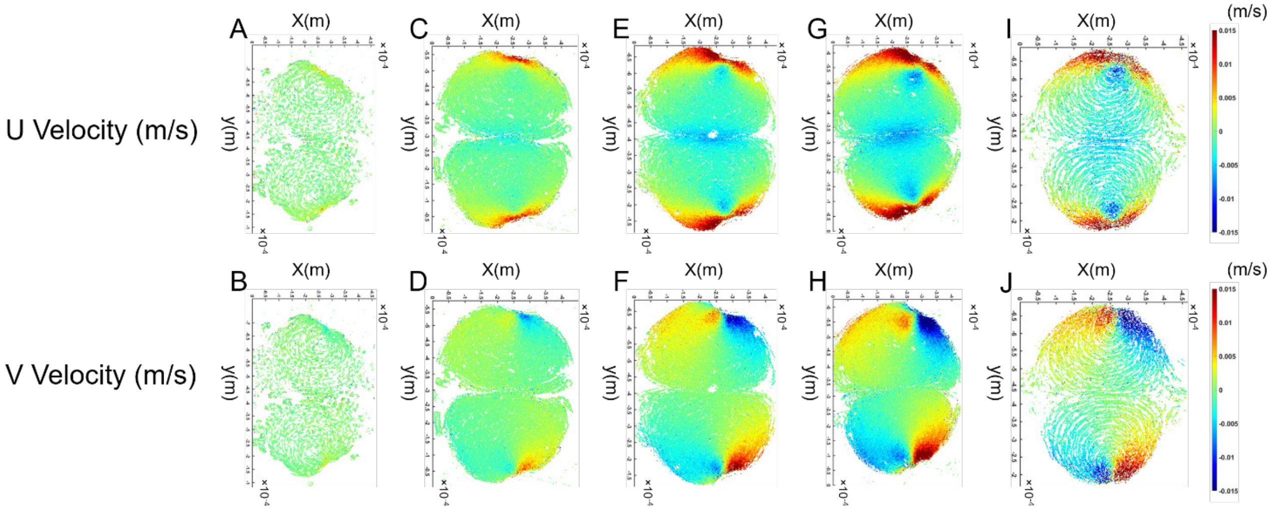


**Figure S1.** PTV analysis results of K562 cells orbiting in acoustic microstreaming vortices at (A&B) PZT voltage=2V, (C&D) PZT voltage=4V, (E&F) PZT voltage=6V, (G&H) PZT voltage=8V, (I&J) PZT voltage=10V

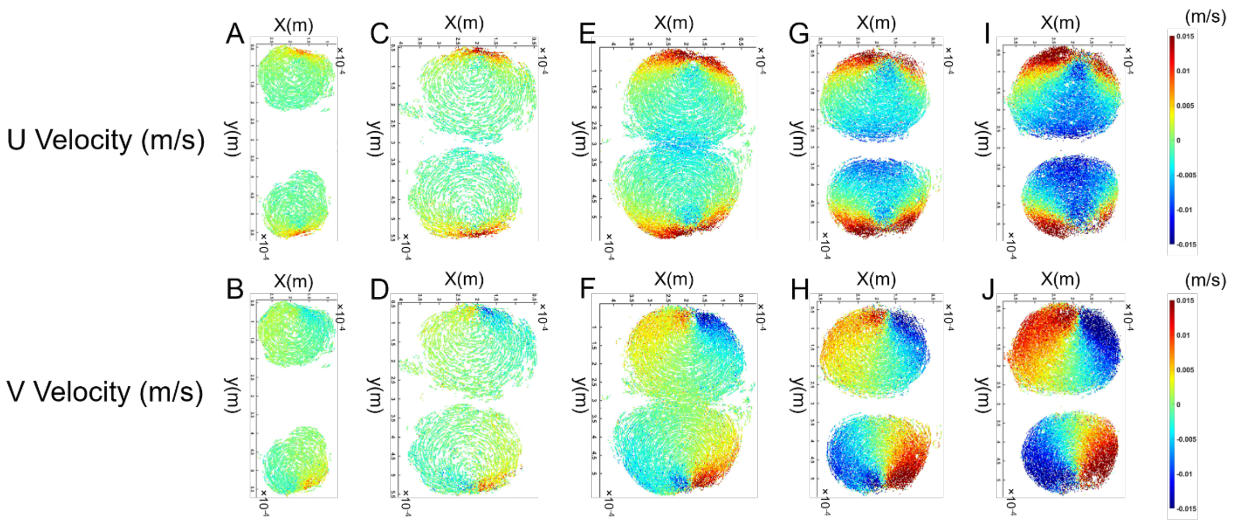


**Figure S2.** PTV analysis results of HeLa cells orbiting in acoustic microstreaming vortices at (A&B) PZT voltage=2V, (C&D) PZT voltage=4V, (E&F) PZT voltage=6V, (G&H) PZT voltage=8V, (I&J) PZT voltage=10V

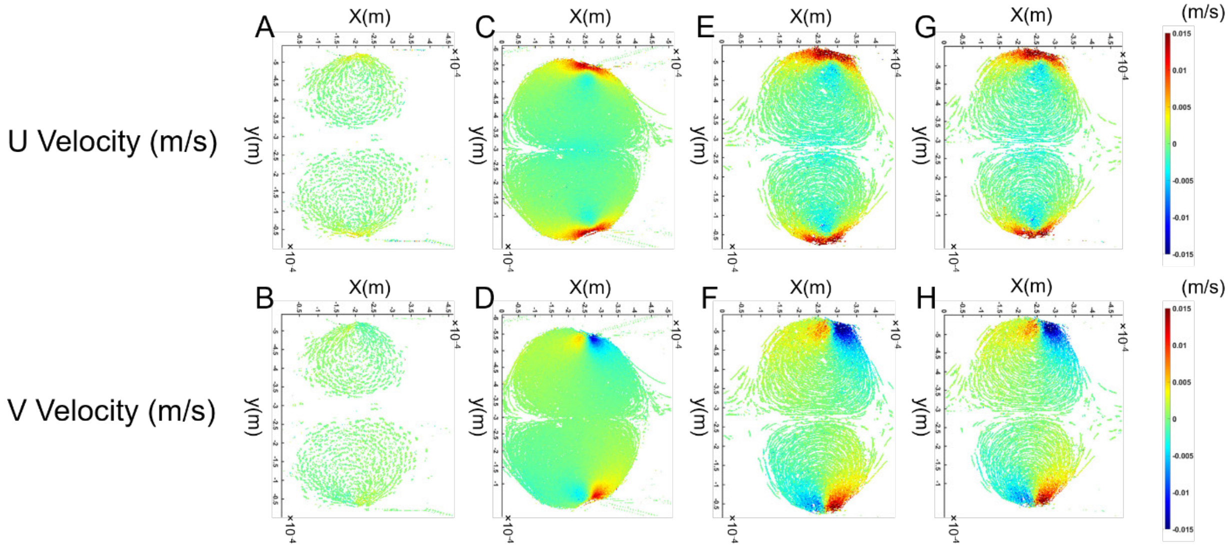


**Figure S3.** PTV analysis results of Jurkat cells orbiting in acoustic microstreaming vortices at (A&B) PZT voltage=2V, (C&D) PZT voltage=4V, (E&F) PZT voltage=6V, (G&H) PZT voltage=10V

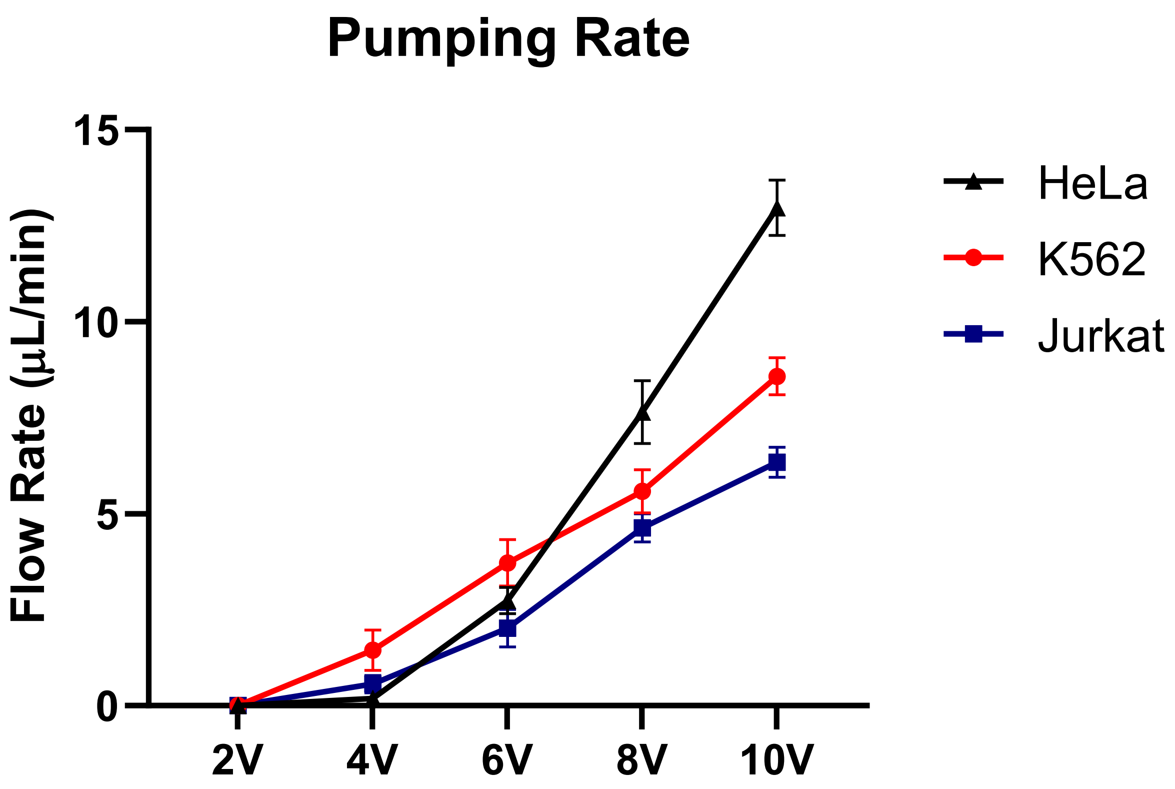


**Figure S4.** Device pumping rate at different PZT applied voltages for HeLa, K562, and Jurkat cells. The unique design of LCATs generates a bulk flow that eliminates the need and complexity of external pumping.

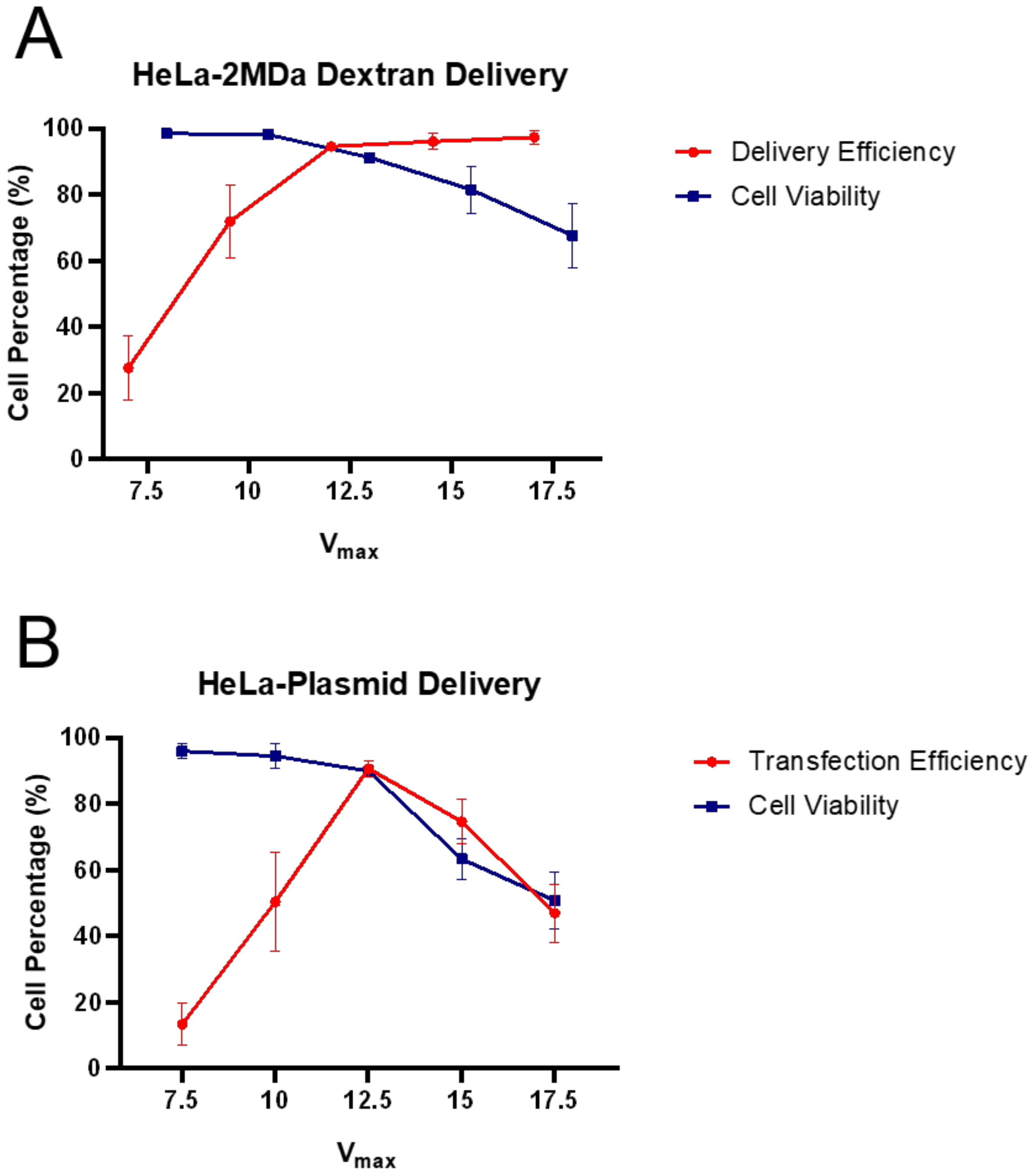


**Figure S5.** Optimization of electric field voltage ($V_{max}$) for **(A)** 2 MDa dextran delivery, and **(B)** eGFP plasmid transfection of HeLa cells at T=10ms and f=10kHz. Based on the results, $V_{max}$=12.5V was found as the optimum applied voltage that resulted in >80% transfection efficiency and >80% cell viability.

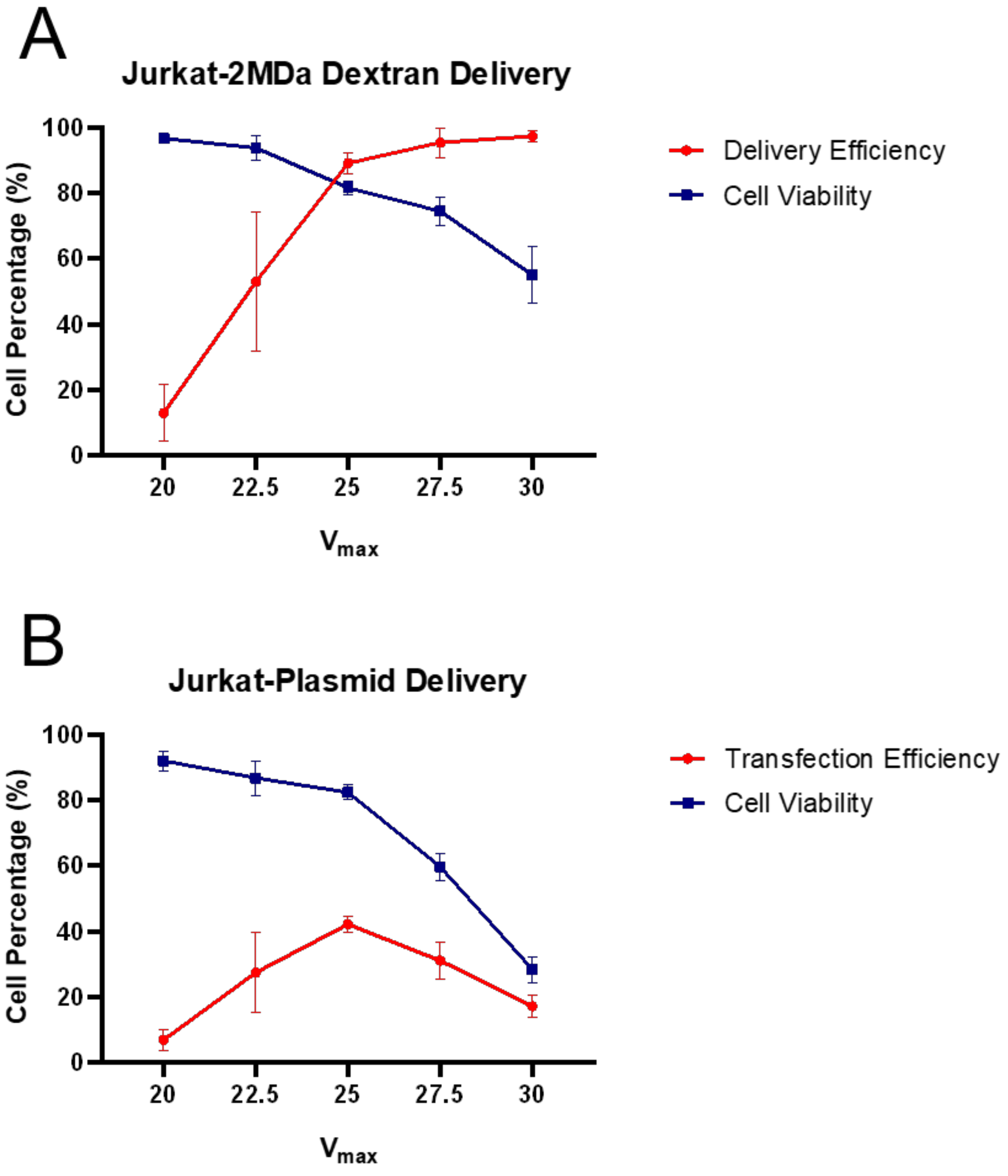


**Figure S6.** Optimization of electric field voltage ($V_{max}$) for **(A)** 2 MDa dextran delivery, and **(B)** eGFP plasmid transfection of Jurkat cells at T=10ms and f=20kHz. Based on the results, $V_{max}$=25V was found as the optimum applied voltage that resulted in >40% transfection efficiency and >80% cell viability (For better demonstration, the data points ±5V of optimum $V_{max}$ are shown).

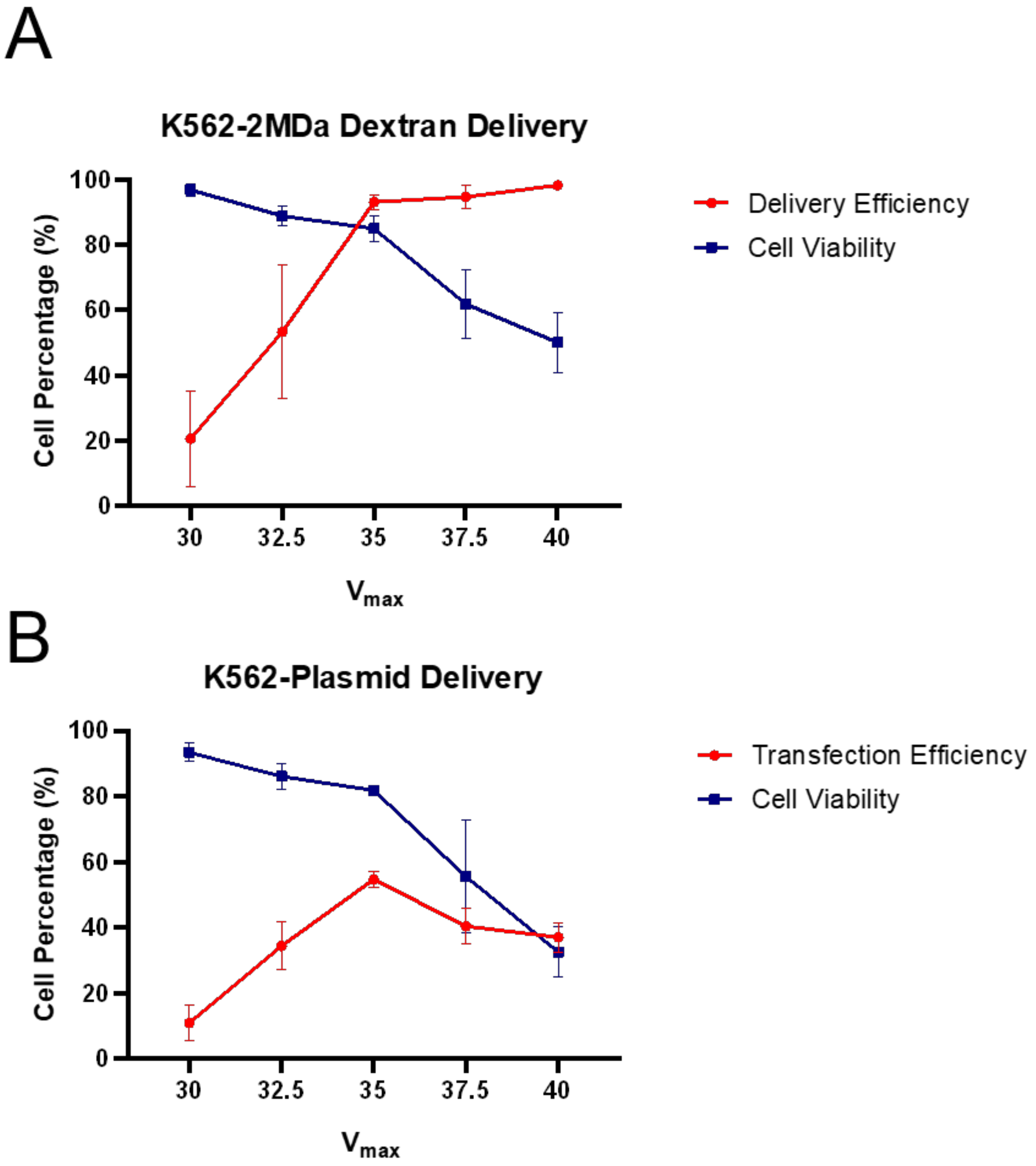


**Figure S7.** Optimization of electric field voltage ($V_{max}$) for **(A)** 2 MDa dextran delivery, and **(B)** eGFP plasmid transfection of K562 cells at T=10ms and f=30kHz. Based on the results, $V_{max}$=35V was found as the optimum applied voltage that resulted in >50% transfection efficiency and >80% cell viability (For better demonstration, the data points ±5V of optimum $V_{max}$ are shown).

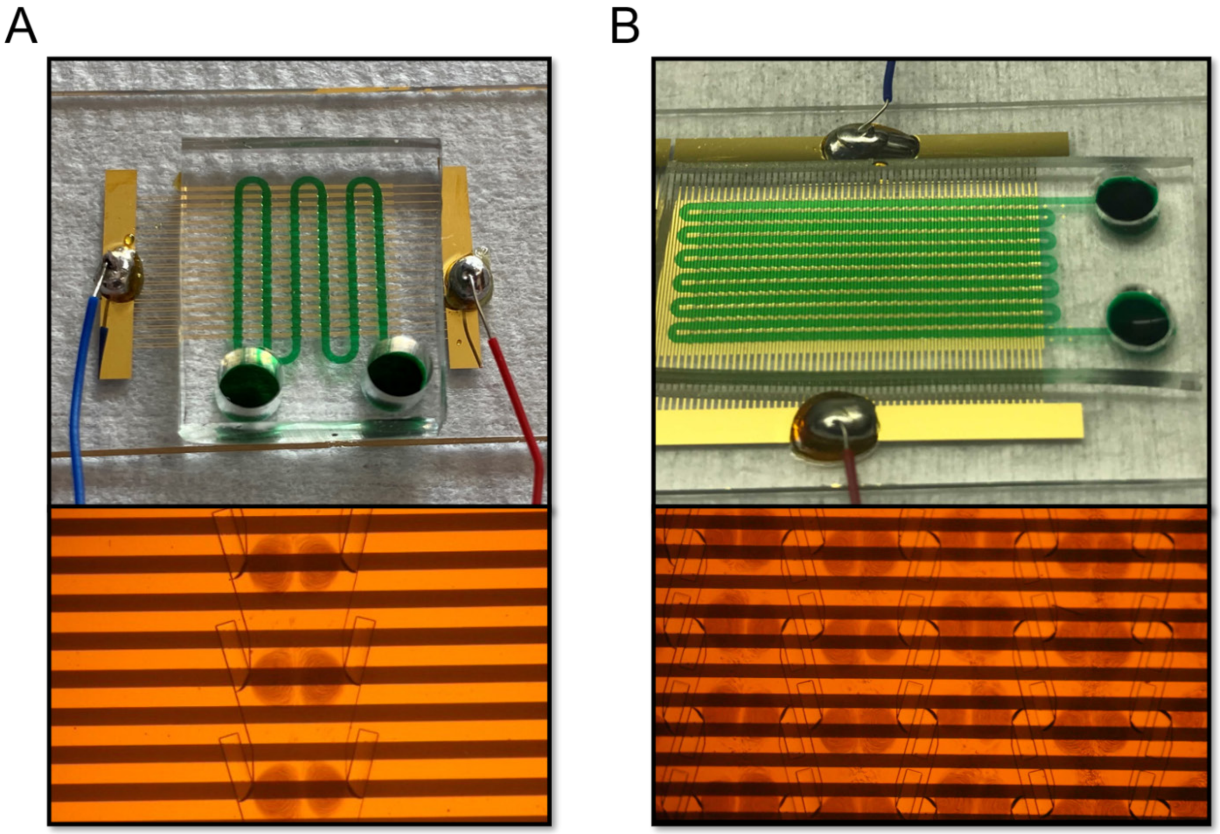


**Figure S8.** Two different AESOP versions. **(A)** Moderate throughput: capable of processing up to 200K cells/min, and **(B)** High throughput: capable of processing up to 1M cells/min

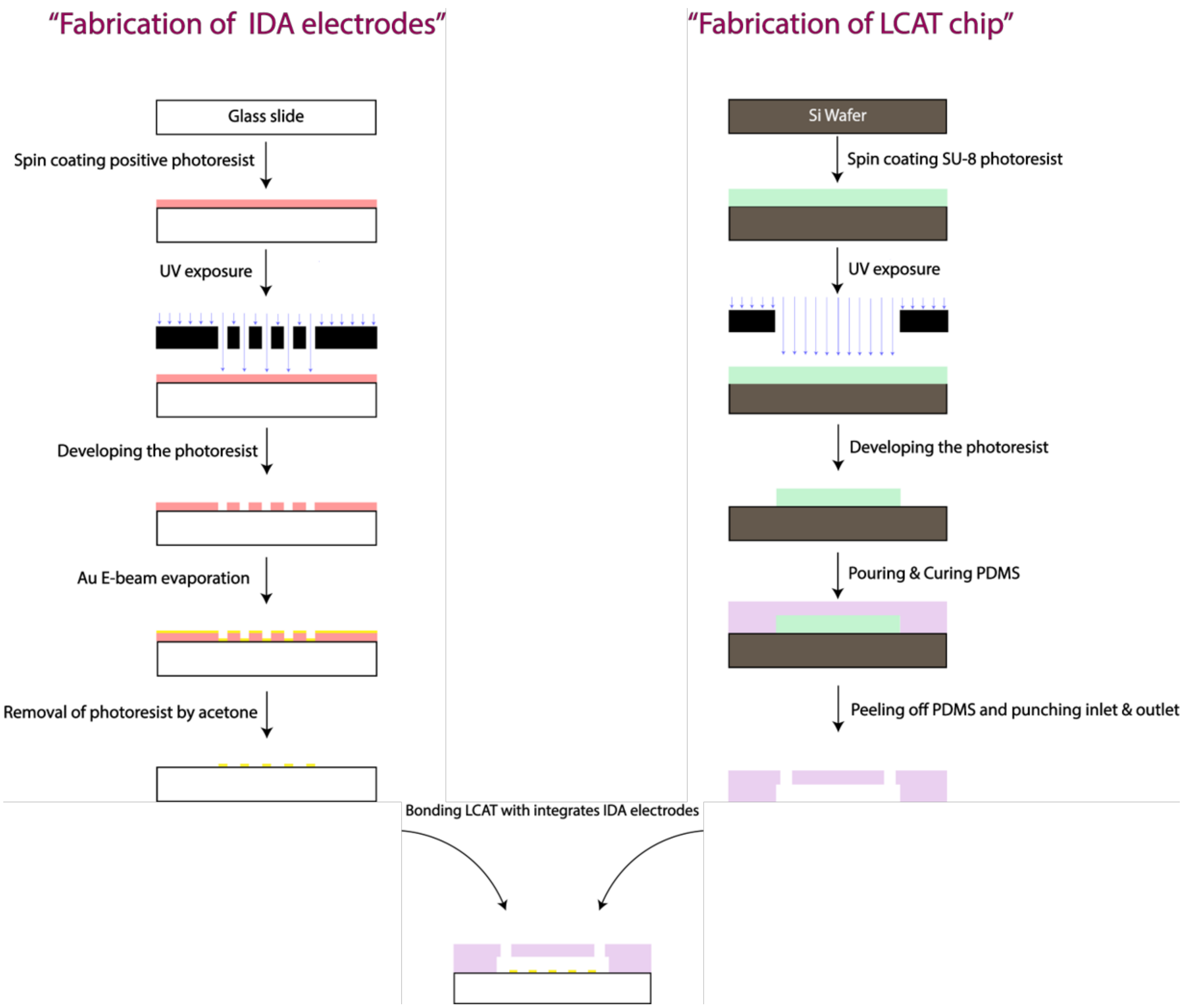


**Figure S9.** Schematics for device fabrication. AESOP integrates interdigitated array (IDA) electrodes with LCAT chip. Lift-off technique was adopted for batch electrode fabrication and soft lithography technique was employed for fabrication of the LCAT chip.
